# Gene-Specific Analysis of Clonal Hematopoiesis Identifies ASXL1 as a Risk Factor for Lung Cancer

**DOI:** 10.64898/2026.05.21.726910

**Authors:** Zijian Zhang, Jing Dong, Yun Huang, Yanhong Liu, Christopher I Amos, Chao Cheng

## Abstract

**Introduction:** Clonal hematopoiesis of indeterminate potential (CHIP) is a recognized risk factor for hematologic malignancies, but its contribution to different types of solid cancers remains incompletely defined.

**Methods:** Here, we performed a systematic, gene-specific analysis of CHIP across 19 common solid cancer types using two large population-based cohorts, the UK Biobank and All of Us with Cox proportional hazards models and nested case-control logistic models.

**Results:** We demonstrate that the relationship between CHIP and solid tumors is highly cancer-type specific, with lung cancer exhibiting the strongest association. In lung cancer, this association is largely driven by ASXL1-mutant clones. Specifically, high variant allele fraction (high-VAF) ASXL1 conferring a significantly increased risk (hazard ratio = 3.2), and the associations remained robust after adjustment for age, sex, body mass index (BMI), smoking status, and genetic ancestry. Notably, ASXL1 CHIP was substantially enriched among smokers, and its association with lung cancer risk was restricted to ever-smokers, highlighting a key interaction between CHIP and environmental exposure. The enrichment of ASXL1 CHIP in lung cancer was further validated in two independent cancer-only cohorts, including MSK-IMPACT and TCGA. In addition, rare germline variant association analysis revealed that germline variation in ASXL1 had the strongest association with lung cancer susceptibility among all solid tumors.

**Conclusions:** Collectively, our findings support a model in which smoking–associated expansion of ASXL1-mutant clones contributes to lung cancer development and suggest that gene-specific CHIP metrics may enhance risk stratification and early detection strategies.

## Introduction

Clonal Hematopoiesis of Indeterminate Potential (CHIP) describes the presence of an expanded hematopoietic stem cell clone carrying somatic mutations in genes commonly implicated in blood cancers, without meeting criteria for a hematologic malignancy^1^. CHIP is an age-related condition and has been consistently associated with increased risks of hematologic neoplasms, chronic inflammation, and cardiovascular disease^2–6^. Although its role in hematologic malignancy is well established, its association with solid cancer risk remains under investigation, and consensus has not been reached^7–10^.

Several population-based studies have investigated the relationship between CHIP and the risk of solid tumors^9–13^. Limited by differences in sample size and heterogeneity in CHIP detection methods, these studies reported modest and sometimes inconsistent associations across multiple solid tumor types^7^. To date, it remains unclear whether CHIP contributes uniformly to the risk of different solid malignancies or exhibits cancer-type–specific effects. Furthermore, the gene-specific effects of CHIP mutations on cancer risk are not well defined, as most studies have treated CHIP as a composite entity rather than examining individual driver genes.

Lung cancer, a common malignancy strongly associated with smoking, has been one of the most extensively studied outcomes in relation to clonal hematopoiesis. Multiple studies have reported a positive association between CHIP and increased lung cancer risk^10–12,14^, including our analyses from the INTEGRAL-ILCCO cohort^15^. For example, Tian et al. demonstrated that this association is largely driven by CHIP mutations with a high variant allele fraction (≥10%)^10^. However, our previous studies suggest that the relationship may be subtype-dependent rather than uniform across the lung cancer patients^16^. Furthermore, the interpretation of these associations is complicated by the presence of therapy-related CHIP mutations, which are frequently observed in patients with solid tumors^17,18^.

In this study, we performed a systematic analysis to investigate the gene-specific effects of CHIP across 19 solid cancer types using two large population-based cohorts, the UK Biobank (UKBB)^19^ and All of Us (AOU)^20^. We demonstrate that the relationship between CHIP and solid tumors is cancer-type specific, with lung cancer showing the strongest association. The association is largely driven by ASXL1-mutant clones with high variant allele frequency (VAF). Importantly, the link between ASXL1 CHIP and lung cancer remains robust after adjusting for established risk factors, including age and smoking status. ASXL1 CHIP is substantially enriched in smokers, and its association with lung cancer risk is confined to this exposure group, with no detectable effect in never-smokers. We further validated the enrichment of ASXL1 CHIP mutations in lung cancer relative to other solid cancers using two independent cancer-only cohorts, MSK-IMPACT and TCGA. Collectively, our analyses support a model in which tobacco smoke–associated expansion of ASXL1-mutant CHIP clones preferentially enriches in lung cancer and contributes to increased disease risk.

## Methods

### Study populations

#### UK Biobank

UK Biobank is a prospective cohort of over 500,000 participants aged 40-70 years between 2006 and 2010 from 22 assessment centers throughout the UK^21^. All participants provided informed consent. Ethical approval for the UKBB was obtained from the Northwest Multi-center Research Ethics Committee. This work uses data provided by patients and collected by the National Health Service as part of their care and support. At enrollment, participants had a physical examination, answered the questionnaire, and provided blood samples. The participants with whole-exome sequencing datasets (N = 469,376) were included in this study. Individuals with prior solid or hematologic malignancies were excluded.

#### All of Us Research Program

The NIH All of Us Research Program is a longitudinal cohort that began enrolling participants aged 18 and older in May 2018 and plans to eventually enroll at least 1 million individuals from the United States^20^. The National Institutes of Health Institutional Review Board approved the study protocol. All participants provided informed consent upon enrollment, including consent to the sample collection and use of their data. This study used data from the Controlled Tier of the Curated Data Repository version 7 (CDR7, release C2022Q4R9), which includes over 413,000 participants. Of these, 245,388 individuals had short-read whole-genome sequencing data available^22^. The individuals with prior cancer diagnoses before the blood collection were excluded.

#### Other cohorts

The MSK-IMPACT^23^ and TCGA^24^ cohorts were used for validation. The MSK-IMAPCT CH cohort includes 39, 510 patients across 23 major cancer types. Patients underwent paired tumor–normal targeted next-generation sequencing as part of clinical genomic profiling at Memorial Sloan Kettering Cancer Center using the MSK⍰IMPACT assay^25^. The TCGA CH cohort contains 2,728 individuals from TCGA who were diagnosed with a first-time primary cancer and no radiation or chemotherapy treatment. The cohort includes patients across 11 diverse cancer types. CHIP mutations were identified from targeted sequencing of matched normal blood samples.

#### Solid cancer ascertainment

In UKBB, solid cancers were identified through national cancer registries using ICD-10 codes^26^. We included 19 of the most prevalent solid cancer types across multiple tissues and organs^27^; the corresponding ICD-10 codes are summarized in the supplementary Table S1. For the patients diagnosed with multiple solid cancer types, the earliest diagnosis of cancer was defined as the primary cancer type. Cases diagnosed before or within 90 days of enrollment were also excluded. Each case was assigned to a single solid cancer type. In the AOU cohort, cancer cases were identified using SNOMED CT codes^28^ (Supplementary Table S1), and the same exclusion criteria as described for the UKBB were applied.

#### Covariates

In UKBB, covariates including age (p21022), sex(p30), ethnic background (p2100), ever/never smoking status (p20160), body mass index (BMI, p21001), date of last personal contact with UKBB (p20143), and principal components of genetic ancestry (p22009) were extracted using the corresponding field IDs. In AOU, demographic data, including date of birth, sex at birth, and race, were retrieved from the PERSON table. The AOU Research Program computed the principal components of genetic ancestry^22^, and they are available on the researcher’s workbench. Smoking status was determined based on responses to the survey question: “Have you smoked at least 100 cigarettes in your entire life?”, categorizing participants into “ever” or “never” smokers.

#### CHIP curation

In UKBB, CHIP mutations were identified from peripheral blood whole-exome sequencing using Mutect2 and filtered according to previously established criteria^29^. CHIP was defined as nonsynonymous or splice-altering mutations in 55 driver genes. 16617 CHIP mutations in 15620 participants were curated. In AOU, similar methods applied to whole-genome sequencing identified 10464 CHIP events in 9493 individuals. In MSK-IMPACT, CHIP variants were identified using Mutect2 and VarDict from target sequencing for the matched normal blood samples, and variants were retained if identified by both callers^24^. In TCGA, whole-exome sequencing data were analyzed using VarScan, GATK, and Pindel with additional filtering criteria applied as previously described^30^. All CHIP findings were defined as the presence of leukemia-associated somatic mutations detected in blood with a variant allele fraction (VAF) ≥ 2^24,29^. Analyses were conducted for overall CHIP and stratified by VAF burden (< 10% vs >= 10%).

#### Statistical analysis

Cox proportional hazards models were fitted using the coxph() function from the R *survival* package. Models were adjusted for age, sex, smoking status, body mass index (BMI), and the principal components of genetic ancestry in the UKBB and AOU cohorts. Follow-up time in the UKBB was calculated from the date of blood collection to the date of first solid cancer diagnosis for incident cases, and to the date of last follow-up or censoring for participants without a cancer diagnosis. In the AOU, follow-up time was similarly defined from the date of blood collection to the date of first solid cancer diagnosis for incident cases. For participants without a cancer diagnosis, follow-up was censored at the most recent electronic health record (EHR) date, representing the last known time the participant was under observation. Nested case-control analyses were conducted within predefined risk windows (10 years for the UKBB and 2.5 years for the AOU). Controls were matched to cases at a 1:5 ratio using nearest-neighbor matching implemented in the *MatchIt* package. Logistic regression models accounting for the matched design were subsequently fitted and adjusted for genetic principal components. All analyses were conducted in R using the cloud platforms provided by DNAnexus or the AOU Research Program.

## Results

### The prevalence of CHIP in UKBB subjects

From UKBB, we selected solid cancer cases representing the 19 most prevalent cancer types (Supplementary Table S2). To minimize potential confounding, we included only individuals who were diagnosed with the corresponding cancer at least 3 months after blood collection (used for WES to identify CHIP mutations) and who had a single cancer diagnosis (i.e., no hematological malignancies or other solid tumors). Among these, prostate cancer was the most prevalent, whereas lung cancer ranked fourth, with 3,034 incident cases (Figure 1A).

**Figure 1:**
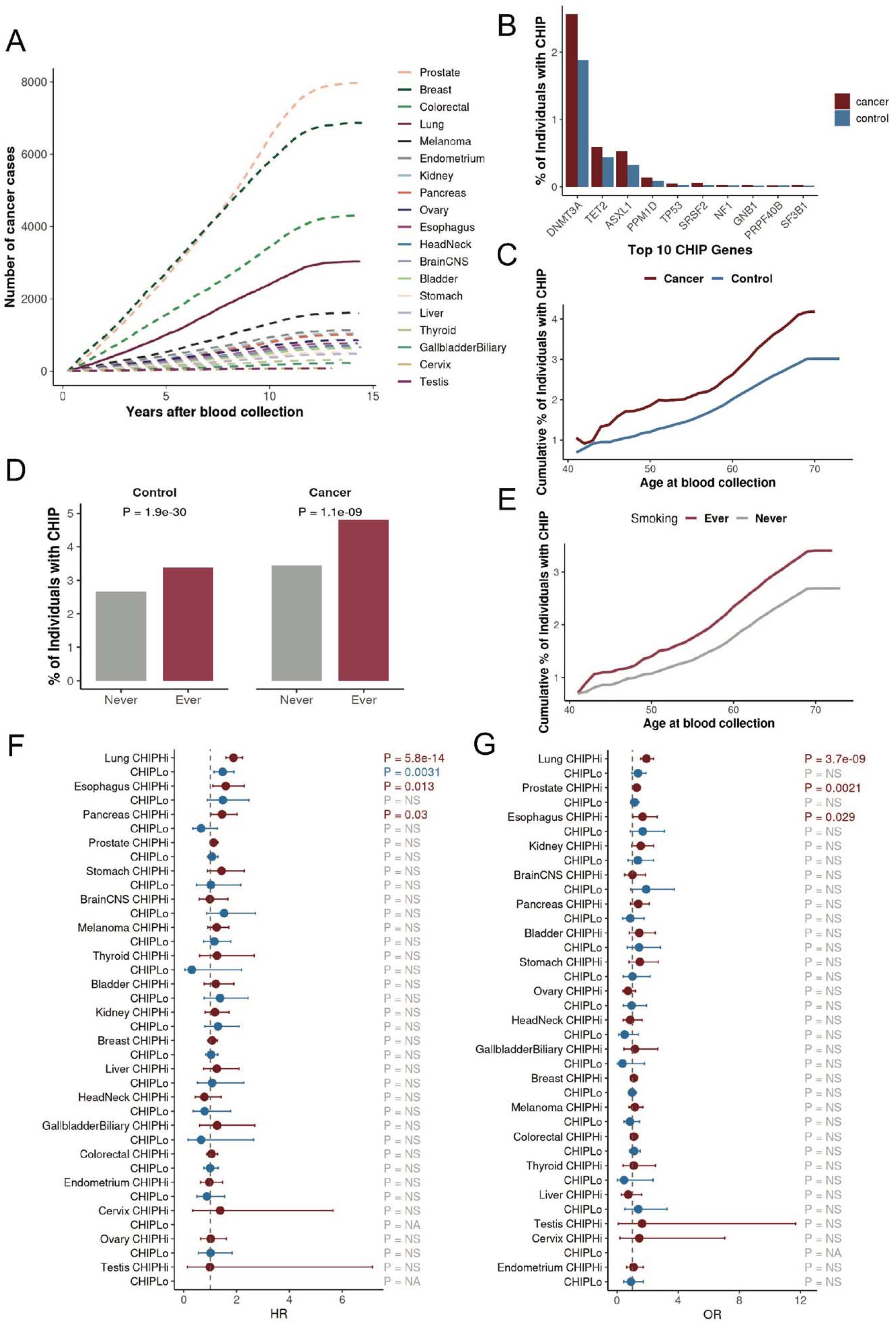
CHIP prevalence and its association with solid cancer risk in the UKBB. (A) Cumulative incidence curves showing the number of incident solid cancer cases following blood collection. (B) Bar plot showing the prevalence of the top 10 most frequently mutated CHIP genes among cancer cases and controls. (C) Cumulative proportion of individuals with CHIP by age at blood collection in participants with solid cancer and controls. (D) Bar plot showing the proportion of individuals with CHIP stratified by smoking status (Ever vs Never) in participants with solid cancer and cancer-free controls. P-values are from chi-square tests. (E) Cumulative proportion of individuals with CHIP by age at blood collection in cancer-free controls, stratified by smoking status. (F) Forest plot showing the association between CHIP and the risk of solid cancers using Cox proportional hazards models. CHIP was categorized as no CHIP, low-VAF CHIP (CHIP-Lo, VAF <10%), or high-VAF CHIP (CHIP-Hi, VAF ≥10%). NS denotes P > 0.05. (G) Forest plot showing the association between CHIP and 10-year risk of solid cancers from nested case-control logistic

In parallel, we identified mutations in 55 CHIP-associated genes among 407574 subjects with WES data, after excluding 8713 individuals with a history of hematologic diseases. Among these subjects, 14,929 (3.7%) carried at least one CHIP mutation. The majority of CHIP+ individuals (93.4%) harbored mutations in a single gene, whereas a smaller fraction had mutations in up to three genes. Across both selected solid cancer cases and non-cancer controls, DNMT3A, TET2, and ASXL1 were the most frequently mutated genes (Figure 1B), consistent with prior large-scale sequencing studies^1,7,24,31^. CHIP prevalence increased with age in both groups and was consistently higher among individuals who developed solid cancer (Figure 1C). Among subjects with solid cancer, 4.2% (1348 of 32322) carried CHIP mutations, which is significantly higher than the 3.0% (9174 of 306,785) observed in non-cancer controls. In both cases and controls, individuals classified as ever-smokers (i.e., current or former smokers), exhibited significantly higher CHIP prevalence compared with never-smokers (Figure 1D). We further examined the age-dependent accumulation of CHIP among non-cancer controls and found that ever-smokers accumulated CHIP mutations at a faster rate than never-smokers before age 60, resulting in a steeper trajectory (Figure 1E); however, this difference diminished after age 60. In contrast, among the cancer cases, CHIP prevalence rose sharply with age and the gap between ever- and never-smokers widened with age. (Supplementary Figure S1). These findings suggest a combined influence of aging and tobacco exposure on clonal expansion.

### Association of CHIP with solid cancer risk

Several previous studies have investigated the association of CHIP with a wide range of diseases in UKBB, including solid cancers^12^. We therefore sought to confirm the association between CHIP and individual solid cancer types using two complementary statistical approaches. First, we calculated the time from blood collection (used for CHIP identification) to the first diagnosis of the corresponding cancer and performed Cox proportional hazards regression analyses to assess associations between CHIP and each solid cancer type. In the cancer-type-specific analysis, the subjects diagnosed with other cancer types were excluded from the cohort. After adjusting for age, sex, body mass index, smoking status, and genetic principal components, CHIP, particularly high-VAF CHIP (≥10%), was associated with increased risk of lung, esophageal, and pancreatic cancers (Figure 1F, Supplementary Table S3). Among these, lung cancer showed the strongest and most statistically significant association. High-VAF CHIP was associated with a 1.9-fold increase in lung cancer risk (HR = 1.9, P = 5.8×10^−1^□), whereas low-VAF CHIP was associated with a more modest but still significant increase in risk (HR = 1.5, P = 0.003).

To further validate these findings, we performed a nested case-control analysis restricted to cancers diagnosed within 10 years after blood collection, selecting non-cancer controls matched on key clinical factors (age, sex, BMI, and smoking status). Results from this analysis were highly consistent with those from the Cox models, again demonstrating the strongest association for lung cancer, particularly among individuals with high-VAF CHIP (Figure 1G, Supplementary Table S4). These results confirmed the previously reported associations between CHIP and lung cancer^10^.

### The CHIP–lung cancer association is primarily driven by ASXL1 mutations

Given its prevalence and strong statistical association with CHIP, we investigated lung cancer in greater detail. Using Cox regression models, we performed gene-specific analyses to evaluate the association of individual CHIP driver genes with incident lung cancer risk while adjusting for key clinical factors. We excluded subjects diagnosed with other cancer types to minimize confounding and restricted the analysis to four CHIP genes with at least 50 mutation carriers to ensure stable effect estimation. Our results indicated that ASXL1 and DNMT3A were associated with a significantly increased risk of lung cancer (Figure 2A, Supplementary Table S5), with high-VAF ASXL1 (HR = 3.2, P = 3×10^−12^) showing a substantially greater risk increase than high-VAF DNMT3A (HR = 1.7, P = 3×10^−^□). These findings were further supported by the nested case-control analysis (Figure 2B). As shown, ASXL1 demonstrated the strongest association, with high-VAF ASXL1 associated with markedly elevated odds of lung cancer (OR = 4.0).

**Figure 2:**
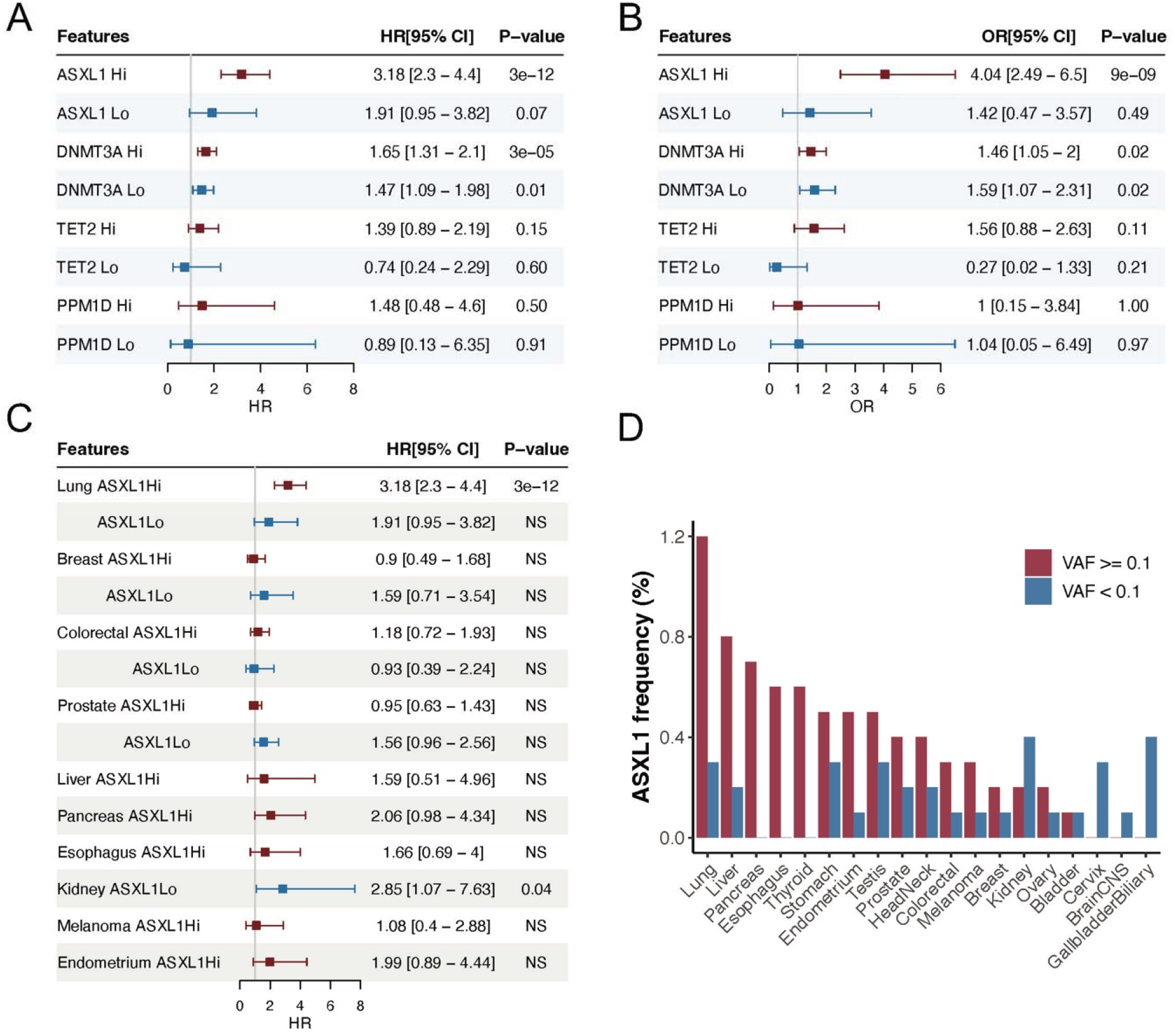
Gene-specific associations between CHIP and lung cancer risk. (A) Forest plot showing the association between CHIP genes and incident lung cancer risk using Cox proportional hazards models. Only CHIP genes with at least 50 carriers are shown. “Hi” and “Lo” denote VAF ≥10% and <10%, respectively, with non-carriers as the reference group. (B) Forest plot showing the association between gene-specific CHIP and incident lung cancer using a time-restricted nested case-control design. (C) Forest plot showing the association between ASXL1 CHIP and incident solid cancer risk using Cox proportional hazards models. Only cancer types with at least three ASXL1- mutant cases in the corresponding group were included. (D) Frequency of ASXL1-mutated individuals across the 19 solid cancer types, stratified by VAF.

The magnitude of these associations prompted us to further investigate whether ASXL1 CHIP is similarly associated with other solid cancer types. Cancer type–specific Cox regression analyses across all 19 solid cancers revealed that ASXL1 CHIP was strongly associated with lung cancer but not with other malignancies (Figure 2C, Supplementary Table S6). Notably, ASXL1 CHIP frequency is highest in lung cancer, especially in high VAF (Figure 2D), further supporting a relationship between ASXL1 clonal burden and increased lung cancer risk. Notably, after excluding all ASXL1 mutations, CHIP remains significantly associated with lung cancer (HR=1.6, P=3×10^-7^, Supplementary Table S7), consistent with a previous report by Tian et al^10^.

Following the identification of a strong link between ASXL1 CHIP and lung cancer risk observed in multivariable Cox regression (Figure 3A, Supplementary Table S8) and nested case-control logistic models (Supplementary Figure S3), the cumulative incidence analysis demonstrated a markedly faster accumulation of lung cancer cases among individuals carrying high-VAF ASXL1 mutations compared with low-VAF carriers and non-carriers (Figure 3B). We next performed stratified analyses to evaluate these associations across different lung cancer histological subtypes. High-VAF ASXL1 was significantly associated with increased risk of lung adenocarcinoma (LUAD, HR = 3.4, P = 2.1×10^−^□) and squamous cell carcinoma (LUSC, HR = 4.1, P = 1.0×10^−^□). In contrast, the association with small cell lung cancer (SCLC) was weaker and only marginally significant (HR = 1.7, P = 0.05), in part reflecting the smaller number of cases for the SCLC histological subtype (Figure 3C, Supplementary Table S9).

**Figure 3:**
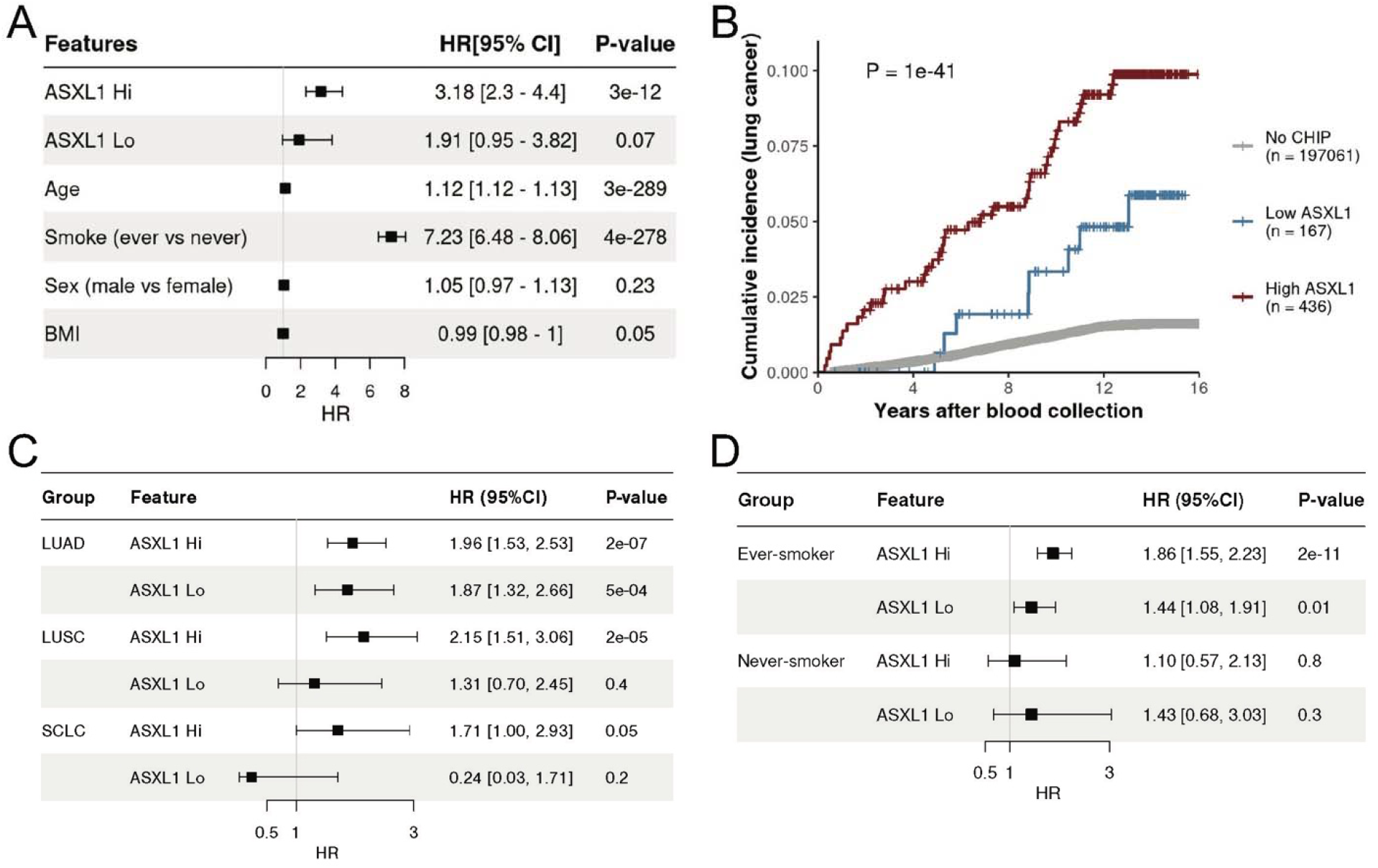
Stratification analysis for ASXL1 CHIP and lung cancer. (A) Forest plot showing the association of ASXL1 CHIP and other covariates with incident lung cancer risk from the Cox proportional hazards model. (B) Cumulative incidence of lung cancer stratified by ASXL1 CHIP status. “Low ASXL1” represents individuals with ASXL1 CHIP and maximum VAF <10%, and “High ASXL1” represents individuals with ASXL1 CHIP and maximum VAF ≥10%. P-value was computed using the log-rank test. (C) Forest plot showing the association of ASXL1 CHIP and lung cancer stratified by the histological subtypes (LUAD: Lung adenocarcinoma; LUSC: Lung squamous cell carcinoma; SCLC: Small cell lung cancer) (D) Forest plot showing the association of ASXL1 CHIP and lung cancer stratified by smoking status (Ever and Never).

Given the well-established link between smoking and lung cancer, we further examined whether the association between ASXL1 CHIP and lung cancer risk differed by smoking status. In analyses stratified by smoking history, high-VAF ASXL1 mutations were significantly associated with increased lung cancer risk among ever-smokers (HR = 1.9, P = 2 × 10^−12^), while low-VAF ASXL1 mutations showed a more modest association (HR = 1.4, P = 0.01). In contrast, no statistically significant association was observed in never-smokers (Figure 3D, Supplementary Figure S3).

### ASXL1 CHIP is enriched in smokers and has the highest mutation rate in lung cancer

The differential association of ASXL1 CHIP with lung cancer risk between ever- and never-smokers may be partly explained by its differential prevalence in these groups. Previous studies have reported that ASXL1 CHIP mutations are more common in smokers^32^. Consistent with this, among lung cancer cases, 46 of 2,776 ever-smokers harbored ASXL1 CHIP mutations, whereas only 2 of 470 never-smokers were carriers (P = 0.02, Fisher’s exact test; Figure 4A). More broadly, in both solid cancer cases and non-cancer controls, ever-smokers were enriched for ASXL1 CHIP mutations compared with never-smokers (Figure 4B). Among lung cancer patients, ASXL1 showed the strongest association with smoking status compared to other CHIP genes (OR = 2.42, P = 1.1 × 10^−^□□; Figure 4C). Furthermore, among ever-smokers, the frequency of ASXL1 CHIP mutations was highest in lung cancer relative to other solid tumor types (Figure 4D). Together, these findings suggest that the pronounced association between ASXL1 CHIP and lung cancer risk is partially driven by its enrichment in smokers. Nevertheless, as noted above, the association between ASXL1 CHIP and lung cancer remains highly significant after adjustment for smoking status, age, and other clinical factors (Figure 3A).

**Figure 4:**
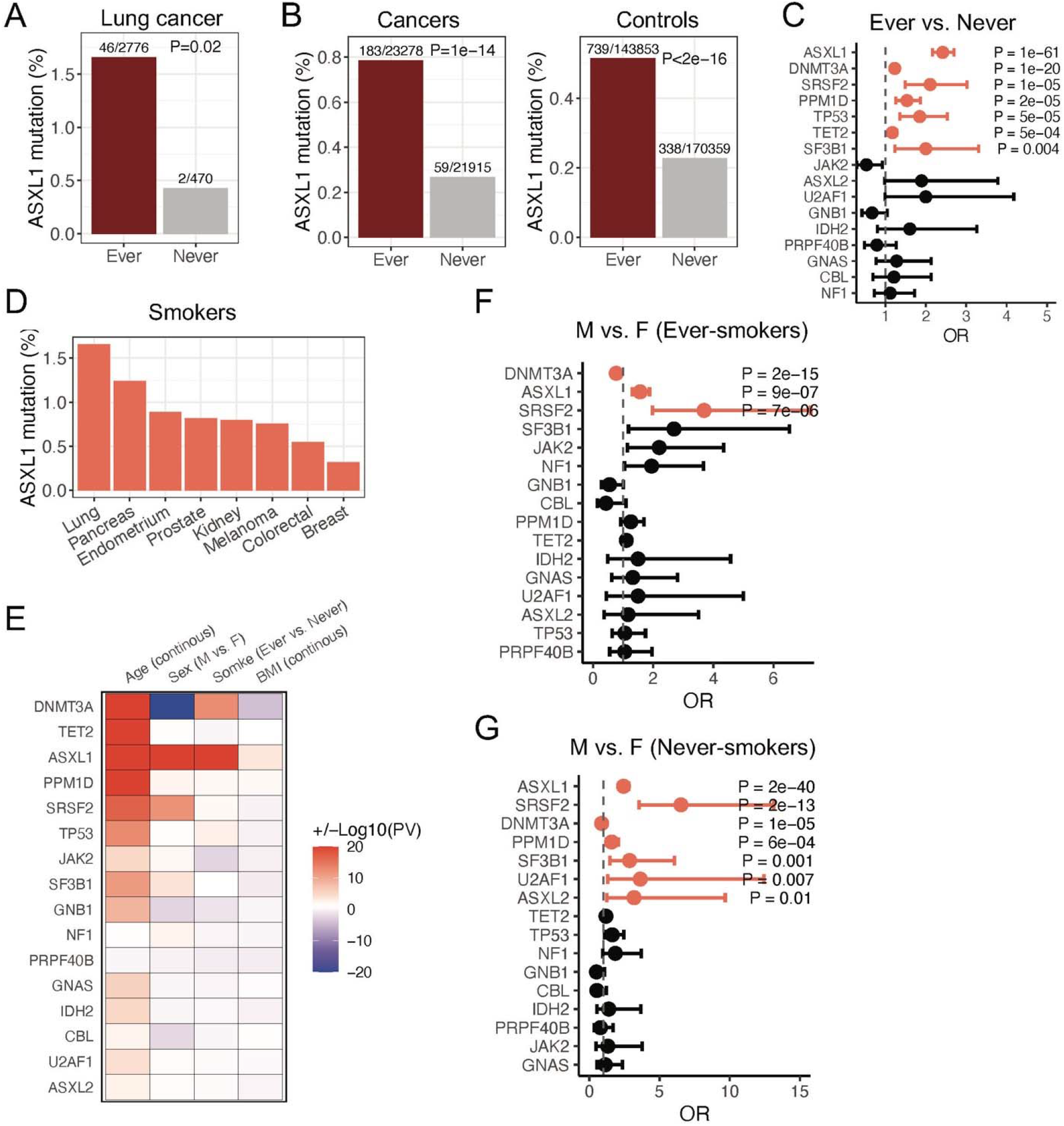
ASXL1 CHIP is enriched in ever smokers. (A) Bar plot showing the percentages of ASXL1 mutated individuals in the ever- and never-smoker groups among lung cancer cases. (B) Bar plot showing the percentages of ASXL1-mutated individuals in the ever- and never-smoker groups across all the cancer cases (left) and controls (right). (C) Forest plot showing the association between CHIP genes and smoking status using logistic regression. Orange color indicates the statistical significance (P < 0.05). (D) Percentage of AXSL1-mutated individuals among smokers with solid cancers. (E) Heatmap showing associations between CHIP genes and age, sex, smoke, and BMI from multivariable logistic regression. Red indicates positive, significant effects; blue indicates negative, significant effects. (F-G) Forest plot showing the stratified analysis of the association between CHIP genes and sex across ever- and never-smokers. Orange denotes statistical significance (P < 0.05).

Furthermore, we used multivariable logistic regression models to investigate the relationships between CHIP and key clinical factors, including age, sex, smoking status, and BMI. Among the most common CHIP genes, ASXL1 was unique in showing significant associations with all of these factors (Figure 4E). After adjustment for other covariates, ASXL1 CHIP was significantly more prevalent in males than in females. Stratified analyses further demonstrated that, in both ever- and never-smokers, ASXL1 exhibited the strongest enrichment in males compared to females among all CHIP genes (Figure 4F-G).

### Validation of the CHIP–lung cancer association in independent Cohorts

In the UKBB data, our systematic analysis revealed a strong association between ASXL1 CHIP and lung cancer. We next sought to validate this finding using the AOU cohort. First, we performed Cox regression analyses to evaluate the association of all CHIP genes with incident lung cancer risk and confirmed that ASXL1 exhibited the strongest association in this cohort (Figure 5A, Supplementary Table S10). Second, we assessed the association of high-VAF ASXL1 CHIP with the risk of common solid cancer types and found that lung cancer was the only solid tumor type showing a significant association (Figure 5B, Supplementary Table S11). As shown, the Cox regression model demonstrated a significant association between high-VAF ASXL1 CHIP and lung cancer risk after adjustment for established clinical factors (HR = 3.4, P = 0.008; Figure 5C), consistent with the findings from UKBB. These results were further supported by nested case-control logistic regression analyses (Supplementary Figure S4).

**Figure 5:**
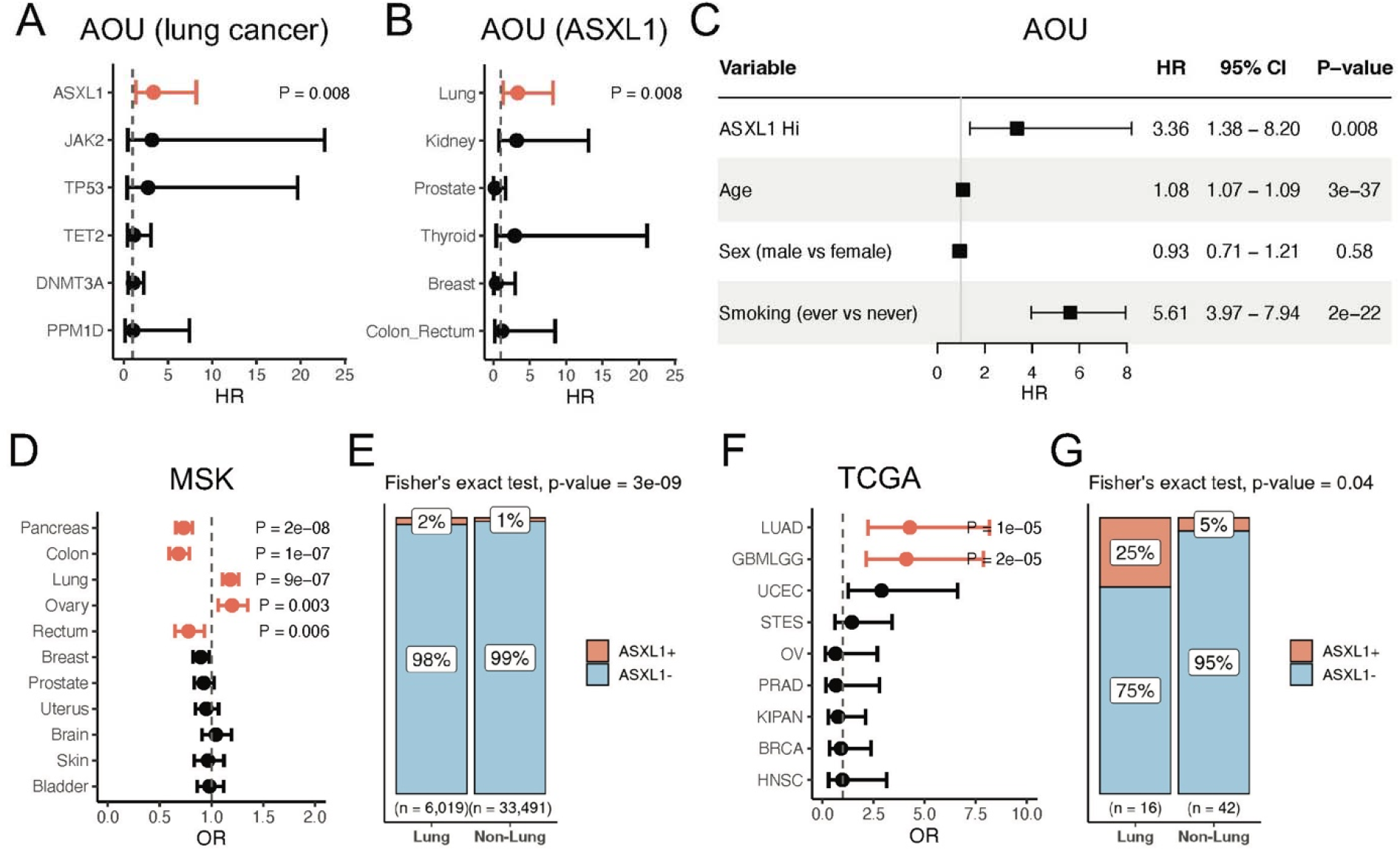
Validation of the association between ASXL1 CHIP and lung cancer risk in All of Us, MSK-IMPACT, and TCGA cohorts. (A) Forest plot showing the association between CHIP genes and incident lung cancer risk in the All of Us (AOU) cohort. Orange denotes statistical significance (P < 0.05). (B) Forest plot showing the association between ASXL1 CHIP and incident solid cancer risk in the AOU cohort. Orange denotes statistical significance (P < 0.05). (C) Forest plot showing the association between ASXL1 CHIP and lung cancer risk in the AOU cohort using the Cox proportional hazards model. Individuals were classified as high ASXL1 (maximum ASXL1 VAF ≥10%), low ASXL1 (maximum ASXL1 VAF <10%), or no CHIP. The ASXL1 low-VAF group was excluded from the forest plot because no lung cancer cases were observed in this category. (D) Forest plot showing the association between ASXL1 CHIP and solid cancer types in the MSK IMPACT cohort (E) Bar plot showing the proportion of patients with ASXL1 CHIP mutations among those with lung cancer versus non-lung solid tumors in the IMPACT cohort. The p-value was calculated using Fisher’s exact test. (F) Forest plot showing the association between ASXL1 CHIP and solid cancer types in the TCGA cohort. (G) Bar plot showing the distribution of ASXL1 CHIP across lung versus non-lung cancer types in TCGA. Statistical enrichment of ASXL1 CHIP in lung cancers compared to non-lung cancers was evaluated using Fisher’s exact test.

Furthermore, we validated the association between ASXL1 CHIP and lung cancer using two additional datasets, MSK-IMPACT and TCGA, which include only solid cancer cases without non-cancer controls. In the MSK-IMPACT cohort, ASXL1 CHIP showed the strongest enrichment in lung cancer compared with other solid tumor types (Figure 5D). As shown, ASXL1 CHIP mutations were significantly enriched among individuals with lung cancer relative to those with non-lung solid tumors (P = 3 × 10^−^□; Figure 5E, Supplementary Table S12). Similarly, in TCGA, ASXL1 CHIP was observed more frequently in patients with lung cancer than in other cancer types (Figure 5F–G, Supplementary Table S13). Taken together, results from UKBB and AOU, as well as these two cancer-only cohorts, demonstrate a robust association between ASXL1 CHIP and increased lung cancer risk.

### Association of ASXL1 with overall survival and other factors

Finally, we examined the gene-specific effects of CHIP on overall survival using Cox regression models, adjusting for age, sex, BMI, and smoking status. When restricted to the UKBB cancer-free controls, we identified six CHIP genes significantly associated with overall survival, all of which were linked to shorter survival (HR > 1; Figure 6A). As shown, TP53 was the most significant CHIP gene (HR = 2.73, P = 5.1 × 10^−^□), while ASXL1 was the second most significant gene (HR = 1.4, P = 5.1 × 10^−^□). A similar survival analysis was performed in a cancer sub-cohort consisting of individuals diagnosed with only one of the 19 selected solid cancer types. In this sub-cohort, three CHIP genes were significantly associated with shorter survival, including ASXL1 (HR = 1.29, P = 0.009), along with SF3B1 and TP53 (Supplementary Figure S5). When individuals were stratified into high-VAF and low-VAF CHIP carriers, only high-VAF ASXL1 was significantly associated with overall survival across all subjects, individuals with solid cancer, and non-cancer controls (Figure 6B). In lung cancer, no significant prognostic association was observed for either high- or low-VAF ASXL1, likely due to limited sample size.

**Figure 6:**
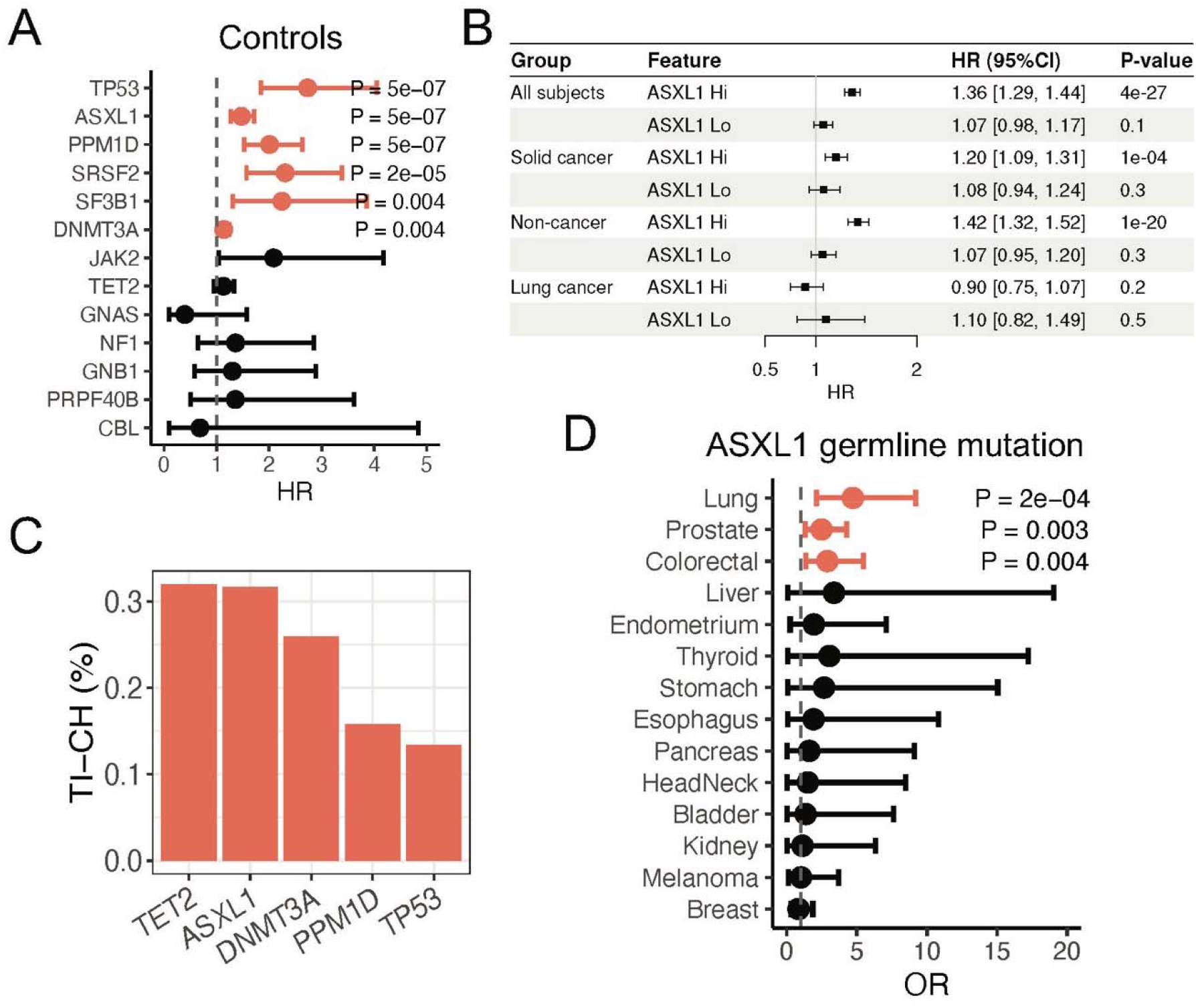
ASXL1 CHIP association with overall survival and tumor microenvironment. (A) Forest plot showing the association of CHIP genes with overall survival across the non-cancer controls in the UKBB. Orange denotes statistical significance (P < 0.05). (B) Forest plot showing the association of ASXL1 CHIP with overall survival across all the subjects, solid cancers, non-cancer controls, and lung cancer cases in the UKBB. (C) Percentage of tumor-infiltrating clonal hematopoiesis (TI-CH) at the gene level among CHIP-positive cancers. TI-CH represents the presence of specific CHIP gene mutations within the tumors, reflecting infiltration of those clones into the tumor tissue by the blood circulation system. (D) Forest plot showing the association between ASXL1 germline variants and solid cancer incidence in the UKBB.

In a pioneering study, Pich et al. evaluated the tumor infiltration clonal hematopoiesis (TI-CH) in non-small cell lung cancer by measuring the presence of CHIP mutations within tumors at a VAF of 2% or more in at least 1 tumor region in the CHIP-positive patients^33^. Although the original study reported TET2 as having the highest rate of TI-CH and therefore focused on TET2 function, reanalysis of their data suggests that ASXL1 shows a comparable TI-CH frequency: TI-CH was detected in 32% of ASXL1 CHIP carriers in lung tumors (Figure 6C). This finding indicates that, similar to TET2, ASXL1 CHIP mutations may also functionally influence the tumor microenvironment in lung cancer^34^.

We have recently systematically identified rare germline variants associated with lung cancer susceptibility and reported that a rare variant in ASXL1 is associated with a significant increase in lung cancer risk^35^. In the UKBB data, we identified a total of 233 individuals carrying the ASXL1 rare germline mutations. Genetic association analysis indicated that individuals with ASXL1 germline mutations had higher susceptibility to lung cancer compared with other solid cancer types (OR = 4.7, P = 2 × 10^−^□). The association between ASXL1 germline mutations and lung cancer risk corroborates our findings from the ASXL1 CHIP analysis and suggests a functional link between this gene and lung cancer susceptibility.

## Discussion

This study systematically evaluated the association between CHIP and the incident risk of 19 common solid cancers in the UKBB. Among these, CHIP, particularly high-VAF clones, was strongly associated with lung cancer. The association remained robust after adjustment for major confounders, including age, sex, smoking status, BMI, and principal components, and was further supported by nested case-control analyses. Previous studies have reported associations between CHIP and lung cancer, though with modest effect sizes^10,11,14,15,36^. By stratifying clones by VAF, our analysis demonstrates that this association is largely driven by high-VAF clones, highlighting the importance of clonal burden in CHIP-related cancer risk. Gene-specific analyses further revealed a particularly strong and specific association between high-VAF ASXL1 mutations and lung cancer incidence, with effect sizes exceeding those previously reported^11,17,36^. This study is among the first to systematically assess CHIP across multiple common solid cancers within a unified framework and to identify one strong gene-specific association with solid cancer risk. These findings highlight gene-specific and VAF-dependent heterogeneity and implicate ASXL1 in CHIP-associated lung cancer risk.

The prominence of lung cancer is biologically plausible, as it shares key risk factors with clonal hematopoiesis, particularly aging and tobacco exposure^2,11,16,37^. Both processes promote somatic mutation accumulation and clonal expansion^38–40^. Experimental models demonstrated that CHIP mutations can actively drive lung cancer progression^17^. For instance, TET2-deficient immune cells in the lung microenvironment produce the inflammatory proteins S100A8 and S100A9, which increase VEGFA secretion and promote tumor angiogenesis^11,17^. Cigarette smoke may contribute to both clonal hematopoiesis in the blood compartment and carcinogenesis in lung tissue, creating a shared etiologic environment linking CHIP to lung cancer development^38–40^.

Among CHIP drivers, ASXL1 showed the strongest association with lung cancer. As an epigenetic regulator and a common CHIP gene^41,42^, germline variants in ASXL1 are associated with increased lung cancer susceptibility^35,43^, supporting the biological plausibility of ASXL1-mediated pathways in lung tumorigenesis^35^. One possible explanation is that chronic inflammation, particularly from tobacco exposure, may promote the expansion of ASXL1-mutant clones^32,40^. Rather than directly inducing ASXL1 mutations, tobacco-related inflammatory stress favors the outgrowth of pre-existing mutant clones^32,38^. These expanded clones may alter immune signaling and hematopoiesis, creating a microenvironment that facilitates lung tumor development^14,17,32^. Thus, tobacco exposure acts primarily as an extrinsic stressor, whereas ASXL1 mutations may contribute independently to lung cancer risk^32,36^.

These findings have potential clinical implications for lung cancer risk stratification. Current screening guidelines primarily rely on age and smoking exposure^44^, which may not fully capture individual susceptibility. High-VAF ASXL1 CHIP could provide additional stratification value and help refine lung cancer screening strategies. Limitations include the potential residual confounding, limited power in some subtype analyses, and the observational design, which precludes causal inference. Future studies are needed to elucidate mechanisms linking CHIP, particularly ASXL1 mutations, to lung carcinogenesis and to determine whether CHIP plays a causal role in reflecting shared risk factors. In conclusion, high-VAF ASXL1 CHIP is strongly associated with incident lung cancer risk, with over a threefold increased risk, underscoring its potential clinical relevance for lung cancer risk stratification and the importance of clonal burden in CHIP-associated cancer susceptibility.

## Abbreviations

CHIP: Clonal hematopoiesis of indeterminate potential
VAF: Variant allele frequency
HR: Hazard ratio
CI: Confidence interval
OR: Odds ratio
UKBB: UK Biobank
AOU: All of Us
MSK-IMPACT: Memorial Sloan Kettering–Integrated Mutation Profiling of Actionable Cancer Targets
TCGA: The Cancer Genome Project
LUAD: Lung adenocarcinoma
LUSC: Lung squamous cell carcinoma
SCLC: Small cell lung cancer;

## Acknowledgements

This research has been conducted using the UK Biobank Resource under Application Number 577088. We thank Dr. Alexander G. Bick and Brian Sharber for providing the CHIP mutation calling set for AOU, which was the foundation for this analysis. We gratefully acknowledge AOU participants for their contributions, without whom this research would not have been possible. We also thank the National Institutes of Health’s All of Us Research Program for making available the participant data, samples, and cohort examined in this study.

## Supplementary Figure and Table Captions

**Supplementary Figure S1:** Cumulative proportion of individuals with CHIP by age at blood collection in cancer-free controls, stratified by smoking status.

**Supplementary Figure S2:** Forest plot showing the association of ASXL1 CHIP and other covariates with incident lung cancer risk from the logistic regression model.

**Supplementary Figure S3:** Forest plot showing the association of ASXL1 CHIP and lung cancer stratified by smoking status (ever and never) using logistic regression models.

**Supplementary Figure S4:** Forest plot showing the association between ASXL1 CHIP and lung cancer risk in the All of Us cohort using the logistic regression model.

**Supplementary Figure S5:** Forest plot showing the association of CHIP genes with overall survival across the cancer cases in the UB biobank.

**Supplementary Table S1:** ICD10/SNOMED codes used to identify solid cancer types and hematological malignancies.

**Supplementary Table S2:** Baseline characteristics of participants with solid cancers and controls in the UK Biobank.

**Supplementary Table S3:** Association between CHIP and solid cancer types in the UK Biobank using Cox regression models.

**Supplementary Table S4:** Association between CHIP and solid cancer types in the UK Biobank using logistic regression models.

**Supplementary Table S5:** Association between individual CHIP driver genes and lung cancer risks using Cox regression models.

**Supplementary Table S6:** Association between ASXL1 CHIP and incident risk across 19 solid cancers using Cox regression models.

**Supplementary Table S7:** Association of CHIP (excluding ASXL1) with lung cancer risk using the Cox regression model.

**Supplementary Table S8:** Association between ASXL1 CHIP and lung cancer risk using the Cox regression model.

**Supplementary Table S9:** Association between ASXL1 CHIP status and lung cancer histological subtypes.

**Supplementary Table S10:** Association between CHIP genes and lung cancer risk in the All of Us cohort.

**Supplementary Table S11:** Association between ASXL1 CHIP status and pan-cancer risk in the All of Us cohort.

**Supplementary Table S12:** Association between ASXL1 CHIP and solid cancer risk (N > 1000) in the MSK-IMPACT CH cohort.

**Supplementary Table S13:** Association between ASXL1 CHIP and solid cancer risk (N > 100) in the TCGA CH cohort.

