## Supplemental Figure S1-5 for "Gene-Specific Analysis of Clonal Hematopoiesis Identifies ASXL1 as a Risk Factor for Lung Cancer"

**Supplementary Figures:**

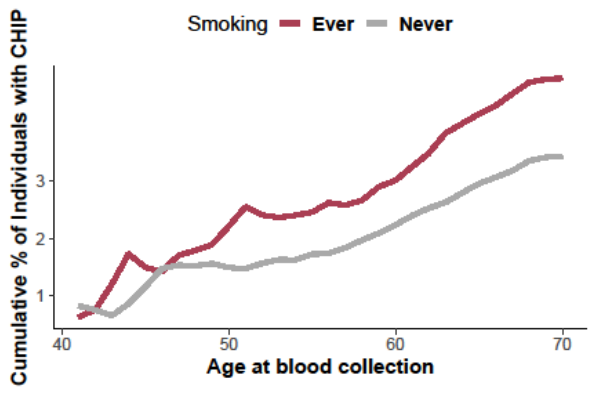

**Supplementary Figure S1:** Cumulative proportion of individuals with CHIP by age at blood collection in cancer-free controls, stratified by smoking status.

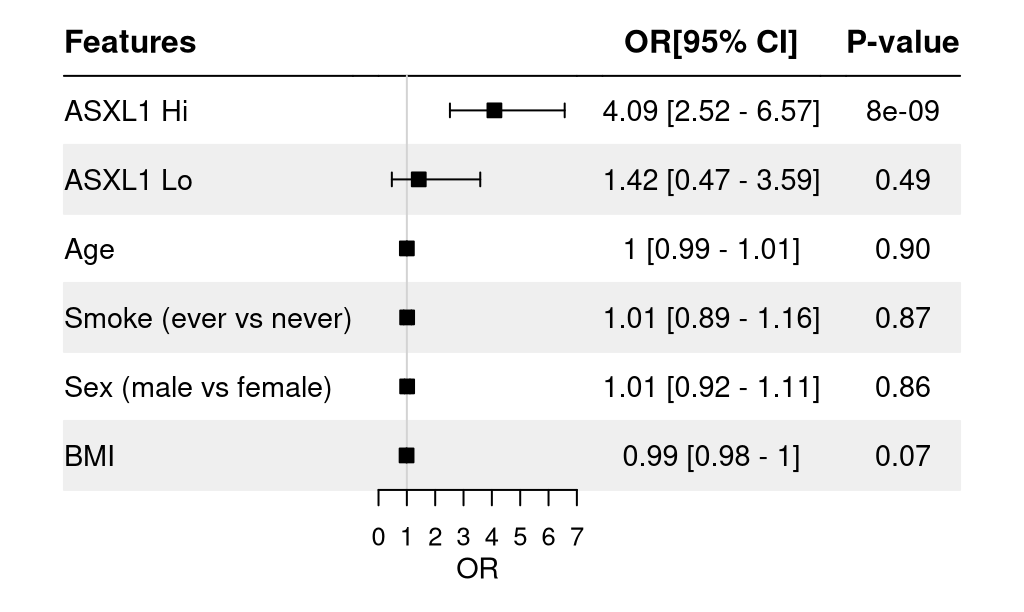

**Supplementary Figure S2:** Forest plot showing the association of ASXL1 CHIP and other covariates with incident lung cancer risk from the logistic regression model.

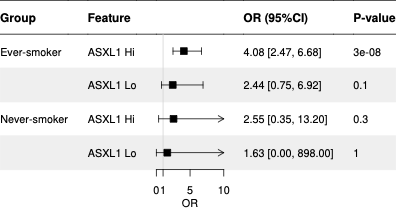

**Supplementary Figure S3:** Forest plot showing the association of ASXL1 CHIP and lung cancer stratified by smoking status (ever and never) using logistic regression models.

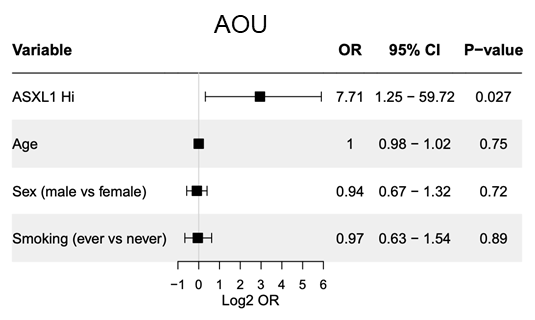

**Supplementary Figure S4:** Forest plot showing the association between ASXL1 CHIP and lung cancer risk in the All of Us cohort using the logistic regression model.

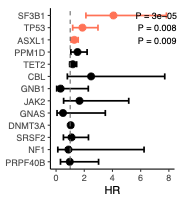

**Supplementary Figure S5:** Forest plot showing the association of CHIP genes with overall survival across the cancer cases in the UB biobank.
